# Different inhibitory interneuron cell classes make distinct contributions to visual perception

**DOI:** 10.1101/275172

**Authors:** Jackson J. Cone, Megan D. Scantlen, Mark H. Histed, John H.R. Maunsell

## Abstract

While recent work has revealed how different inhibitory interneurons influence cortical responses to sensory stimuli, little is known about how their activity contributes to sensory perception. Here, we optogenetically stimulated different genetically defined interneurons (parvalbumin (PV), somatostatin (SST), vasoactive intestinal peptide (VIP)) in visual cortex (V1) of mice working at threshold in contrast increment or decrement detection tasks. The visual stimulus was paired with optogenetic stimulation to assess how enhancing V1 inhibitory neuron activity synchronously during cortical responses altered task performance. PV or SST activation impaired, while VIP stimulation improved, contrast increment detection. Notably, PV or SST stimulation also impaired contrast decrement detection, when opsin-evoked inhibition would exaggerate stimulus-evoked decrements in firing rate, and thus might improve performance. The impairment produced by PV or SST stimulation persisted throughout many weeks of testing. In contrast mice learned to reliably detect VIP activation in the absence of natural visual stimulation. Thus, different inhibitory signals make distinct contributions to visual contrast perception.

## Introduction

In the visual system, local inhibitory interneurons shape fundamental sensory computations like stimulus tuning (Liu et al., 2011; Priebe and Ferster, 2008), response normalization (Carandini and Heeger, 2012), and center-surround suppression (Adesnik et al., 2012; Ferster, 1988). This wide-ranging influence depends in part on their genetic, physiological, and morphological diversity (Markram et al., 2004). Transgenic mice provide selective access to three genetically defined interneuron classes that express parvalbumin (PV), somatostatin (SST), or vasoactive intestinal peptide (VIP). These three classes encompass the vast majority of inhibitory neurons in the cerebral cortex, although each is likely to include distinct inhibitory cell types that differ along many dimensions, especially morphological (Rudy et al., 2011).

Optogenetic tools targeting PV, SST or VIP neurons in mice have helped clarify how different sources of inhibition augment stimulus-evoked responses in sensory cortices. PV and SST neurons, which target somata and dendrites respectively (Kubota et al., 2016; Rudy et al., 2011), suppress visual responses in pyramidal output cells (Atallah et al., 2012; Cottam et al., 2013; Glickfeld et al., 2013), although it remains uncertain whether PV and SST neurons produce quantitatively similar effects on pyramidal cell output (Atallah et al., 2012; Lee et al., 2012; Seybold et al., 2015; Wilson et al., 2012). In contrast, VIP neurons inhibit other inhibitory neurons (Pfeffer et al., 2013; Pi et al., 2013), and increase the gain of pyramidal cell sensory responses (Fu et al., 2014; Pi et al., 2013). Given that VIP neurons gate sensory responses, VIP neurons may enhance sensory perception. However, relatively little is known about how different sources of inhibition influence sensory perception.

Perturbing PV neurons during perceptual tasks has produced conflicting results: some show that PV stimulation improves perceptual ability (Aizenberg et al., 2015; Lee et al., 2012), while others report impaired performance (Glickfeld et al., 2013; Guo et al. 2014). In these studies, optogenetic stimulation was always delivered before, during, and after the target stimulus, leaving uncertainty about how dynamic inhibitory signals contribute to the perception of specific sensory events. Finally, how the activity of SST or VIP neurons augments perception remains largely uninvestigated. Determining the contributions of different inhibitory cell classes is central to understanding the circuits and computations that elaborate perception in sensory cortex.

Here, we expressed the excitatory opsin Channelrhodopsin-2 (ChR2; Nagel et al., 2003) in either PV, SST or VIP interneurons in mouse visual cortex (V1). Mice were highly trained, such that we could precisely measure their behavioral thresholds for detecting brief increments or decrements in visual contrast. We optogenetically potentiated V1 interneuron activity synchronously with visual stimulus presentation to determine how each interneuron class influenced task performance. Our results demonstrate that different inhibitory neurons make distinct contributions to contrast perception.

## Results

Our primary goal was to determine how different classes of inhibitory neurons contribute to visual contrast perception. We used transgenic mouse lines that express Cre-recombinase selectively in one of the three major genetic subclasses of inhibitory neurons in mouse cerebral cortex (PV, JAX strain #017320; SST, JAX strain #013044; and VIP, JAX strain #010908; Hippenmeyer et al., 2005; Taniguchi et al., 2011). Each mouse was surgically implanted with a headpost and a cranial window over cortex to give stable optical access to V1 (Goldey et al., 2014). Following implantation, we localized V1 using intrinsic signal imaging (Figure 1A) to guide injections of a Cre-dependent virus containing ChR2-tdTomato (Figures 1B-C; Nagel et al., 2003). Once ChR2 expression could be verified through the cranial window (∼2 weeks post injection), mice were trained to perform a contrast increment detection task while head fixed. In this task (Figure 1D), head fixed mice faced a video display filled with a mid-level gray. To start a trial, mice depressed and held a lever through an unpredictable delay period (400-3000ms). Following the onset of a brief stimulus (100 ms), mice were required to release the lever within a short response window (700-900 ms) to receive a reward. We used static, vertically-oriented, monochromatic, odd-symmetric Gabor patches (centered at 25° azimuth, −15° to 15° elevation, 6.75° SD; spatial frequency 0.1 cycles/degree). False alarms and missed stimuli resulted in a brief time out (2 s) before the start of the next trial. We varied the visual stimulus contrast to measure psychophysical detection threshold. Visual stimuli were presented at a visual field location corresponding to the V1 representation with strongest ChR2 expression (Figure 1B). An optical fiber was permanently mounted to the headpost to ensure optogenetic manipulations would affect neuronal activity in a consistent location each day.

**Figure 1.**
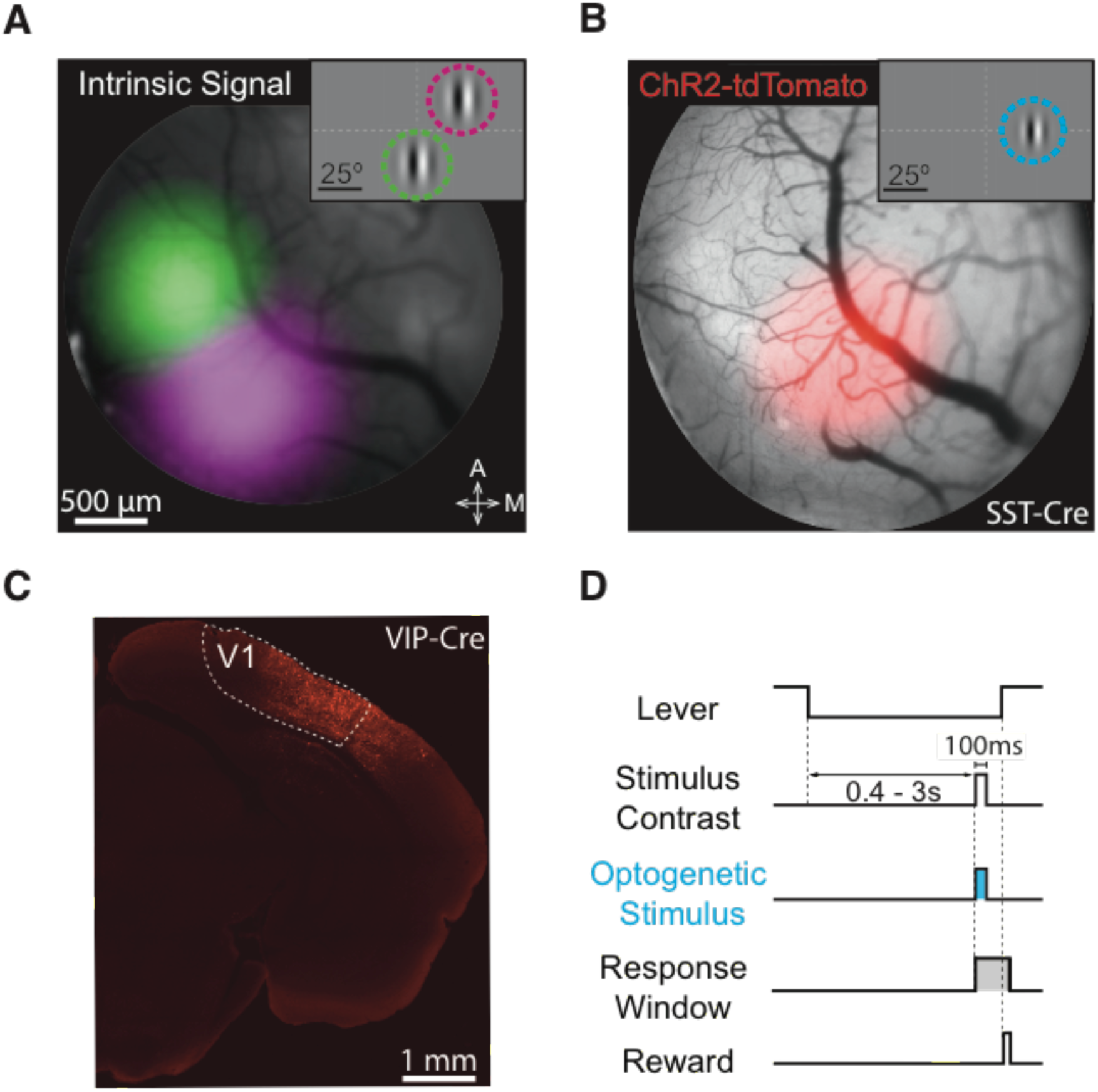
Targeting ChR2 to retinotopically defined areas of visual cortex. (A) Intrinsic autofluorescence responses to visual stimuli presented in two locations in a SST-Cre mouse. Magenta and green features represent 2D-gaussian fits of responses to stimuli at the monitor locations depicted in the inset (green: 0° azimuth, −20° elevation; magenta: 25° azimuth, +20° elevation; Gabor SD = 10°). Dashed lines represent horizontal and vertical meridians. A: anterior; M: medial. (B) ChR2-tdTomato fluorescence (2D-gaussian fit) from the same cortical region shown in A. Expression is maximal at the retinotopic location used in all behavioral sessions (inset; 25° azimuth, 0° elevation; Gabor SD = 6.75°). Cyan ring represents retinotopic location of ChR2 stimulation. Conventions are the same as in A. (C) Representative confocal image of ChR2-tdTomato expression in the visual cortex of a VIP-Cre mouse (D). Trial schematic of the contrast increment detection task. Following the intertrial interval, a trial begins when the mouse depresses the lever. A visual stimulus can appear from 400 to 3000 ms following trial onset. The mouse must release the lever within 700-900 ms after stimulus onset to receive reward. On a randomly selected half of the trials, ChR2-expressing interneurons were illuminated with blue light for 100 ms concurrent with the visual stimulus.

### Visual detection is impaired by PV or SST neuron activation and enhanced by VIP neuron activation

We began optogenetic stimulation experiments as soon as contrast increment detection thresholds were stable. We optogenetically stimulated ChR2 expressing inhibitory interneurons on a randomly selected half of trials. Optogenetic stimulation was delivered concurrently with the visual stimulus (100 ms duration beginning at visual stimulus onset; Figure 1D). The optogenetic stimulus power was held constant throughout a session at a level chosen to produce ∼two-fold change in threshold (typically 0.5–2.0 mW depending on genotype and ChR2 expression). We quantified the effects of optogenetic stimulation by fitting a sigmoid function to behavioral data collected with or without optogenetic stimulation (Figure 2). There were no significant threshold differences detected across genotypes in the absence of optogenetic perturbations (PV median threshold 6.0%; SST median threshold 6.6%; VIP median threshold 6.5%; p = 0.73; Kruskal-Wallis Test).

**Figure 2.**
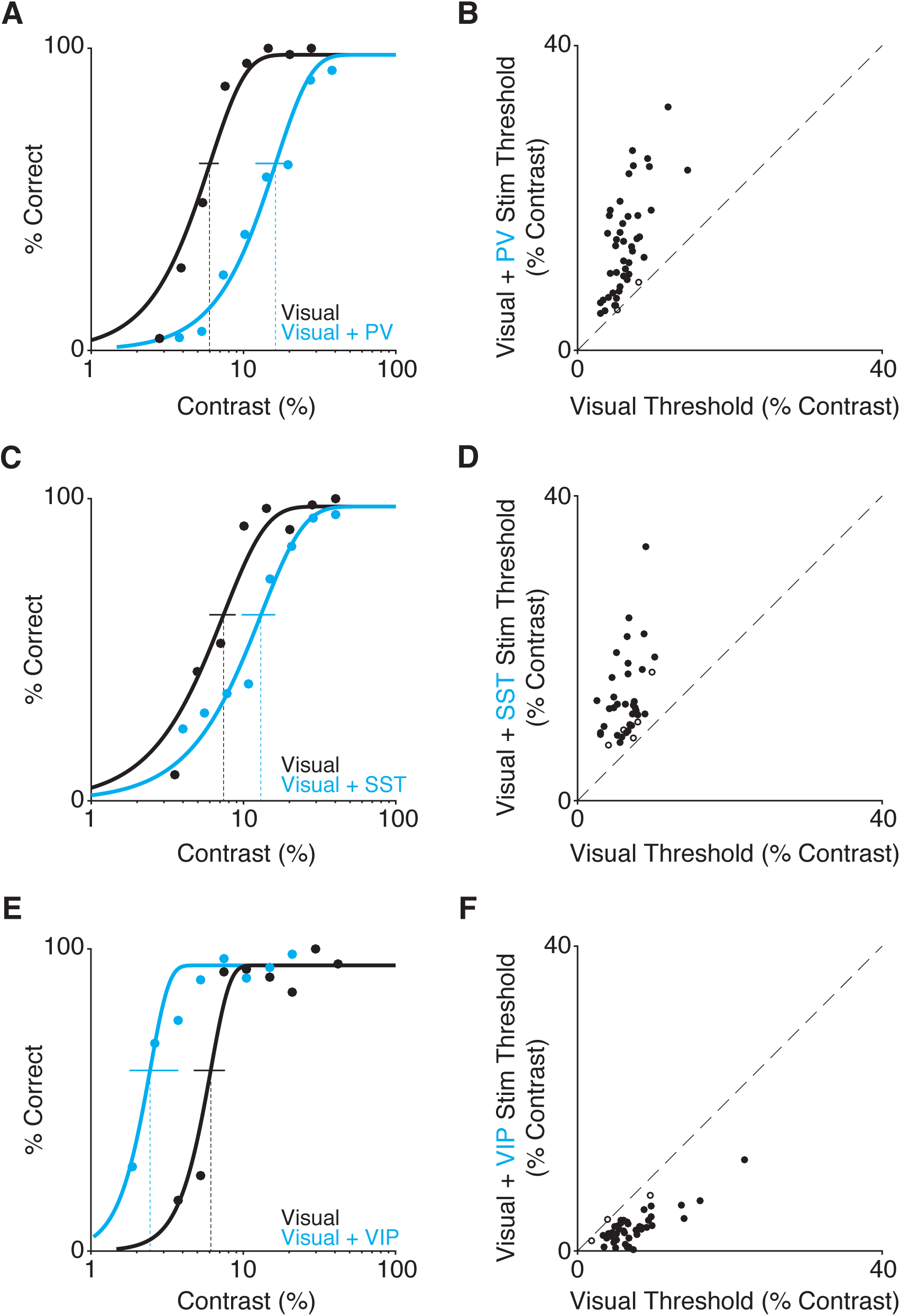
PV and SST stimulation impairs, while VIP stimulation improves, contrast increment detection. (A) Representative PV mouse behavioral performance from a single contrast increment session. Filled dots represent false-alarm corrected performance for trials with (blue) and without (black) activation of PV interneurons as stimulus contrast is varied. Curves are best-fitting Weibull functions that were used to determine detection thresholds (dotted vertical lines) and 95% confidence intervals (solid horizontal lines). (B) Summary of PV stimulation effects. Circles depict the contrast increment detection thresholds from an individual session with (y-axis) and without (x-axis) PV neuron stimulation (5 mice, 47 sessions). Filled circles indicate a significant shift in threshold (44/47; bootstrap). (C) Representative behavioral performance from a single session in a SST mouse. Same format as A. (D) Summary of SST stimulation effects (3 mice, 40 sessions; significant threshold difference in 34/40 sessions). Same format as B. (E) Representative single session behavioral performance from a VIP mouse. Conventions as in A and C. (F) Summary of VIP stimulation effects (3 mice 43 sessions; significant threshold difference in 40/43 sessions). Same format as B and D.

Optogenetic stimulation of PV neurons impaired contrast increment detection, resulting in rightward shift in the psychometric function (Figure 2A). Across all PV mice (5 mice, 47 sessions), contrast increment detection thresholds were significantly higher with PV stimulation (Figure 2B, median 13.6% versus 6.0%, p < 10^−8^; Wilcoxon signed rank test). We expressed the effect on performance as the ratio of thresholds with and without optogenetic input. Values greater than one indicate that optogenetic stimulation impaired detection. On average, PV stimulation produced about a two-fold increase in detection threshold (2.2, SEM 0.1; range 1.0–4.3), although it should be noted that the magnitude of the change depends on factors such as the power of the optogenetic illumination, the level of ChR2 expression and the precise positioning of the optic fiber. Using a bootstrap analysis (see Methods) we found that PV stimulation significantly impaired detection on most of the daily sessions (44/47, Figure 2B).

Optogenetic stimulation of SST interneurons similarly impaired contrast increment detection, shifting psychometric functions to the right (Figure 2C). Across all SST stimulation sessions (3 mice, 40 sessions), contrast increment detection thresholds were significantly greater when the visual stimulus was paired with SST activation (Figure 2D, median 12.0% versus 6.6, p < 10^−7^; Wilcoxon signed rank test). SST stimulation also produced about a two-fold increase in detection thresholds (2.2, SEM 0.1; range = 1.1–5.1). SST stimulation significantly impaired detection on most of the daily sessions (34/40). Overall, enhancing either PV or SST interneuron mediated inhibition similarly impaired the detection of contrast increments.

In stark contrast to the results of PV and SST perturbations, optogenetic stimulation of VIP neurons enhanced contrast increment detection, shifting psychometric functions to the left (Figure 2E). Across all VIP sessions (3 mice, 43 sessions), contrast increment detection thresholds were significantly lower in the presence of VIP stimulation (Figure 2F, median 2.8% versus 6.5, p < 10^−7^; Wilcoxon signed rank test). The effect of VIP activation was about two-fold, but unlike PV and SST stimulation, it was a halving of detection thresholds (0.4, SEM 0.03; range = 0.02-1.0 VIP stimulation significantly improved detection on most daily sessions (40/43). Thus, VIP interneurons make a contribution to visual perception that is the inverse of that made by PV and SST interneurons.

### Contrast decrement detection is impaired by PV or SST neuron activation

A straightforward strategy for detecting contrast increments is to integrate spikes from sensory cortex and respond once the spike rate exceeds a criterion value (Geisler, 2011). Thus, our results in the increment detection task might be explained by how each interneuron type influences pyramidal cell spiking. Pyramidal cell responses to sensory stimuli are increased by VIP activation (Pi et al., 2013), and decreased by PV or SST activation (Atallah et al., 2012; Cottam et al., 2013). Thus, the addition or subtraction of stimulus-evoked spikes in V1 could accelerate (VIP) or impede (PV/SST) progress toward a detection criterion. Conversely, decrements in stimulus contrast might be expected to decrease the firing rate of many pyramidal cells. Therefore in a contrast decrement task, PV or SST stimulation might improve performance because opsin-evoked inhibition would exaggerate the decrement in firing rate produced by the visual stimulus.

To examine this, we retrained a subset of PV and SST (2 of each) on a contrast decrement detection task. A 75% contrast stimulus was always present on the video display except when its contrast transiently decreased to a lower value (100 ms, Figure 3A). Without optogenetic perturbations, there were no differences in contrast decrement detection performance (PV median threshold 10.5%; SST median threshold 11.0%; p = 0.89; Wilcoxon rank-sum test). Surprisingly, PV or SST activation during contrast decrements impaired detection, shifting the psychometric functions to the right (Figure 3B; Figure S1), just as it did for contrast increment detection. Across all sessions (4 mice, 38 sessions), contrast decrement detection thresholds were significantly greater when the visual stimulus was paired with either PV or SST activation (Figure 3C, median = 22.8% versus 10.7, p < 10^−7^; Wilcoxon signed rank test). This effect was significant for both gentoypes (PV median = 23.2% versus 10.5, p < 10^−4^; SST median = 21.4% versus 11.0, p < 10^−3^; Wilcoxon signed rank tests). PV or SST stimulation resulted in an approximately two-fold increase in detection thresholds (PV 2.41, SEM 0.1, range 1.3–4.2; SST 2.0, SEM 0.1, range 1.3–3.0). A significant increase in detection threshold was found in most individual sessions (PV: 20/20; SST 17/18).

**Figure 3.**
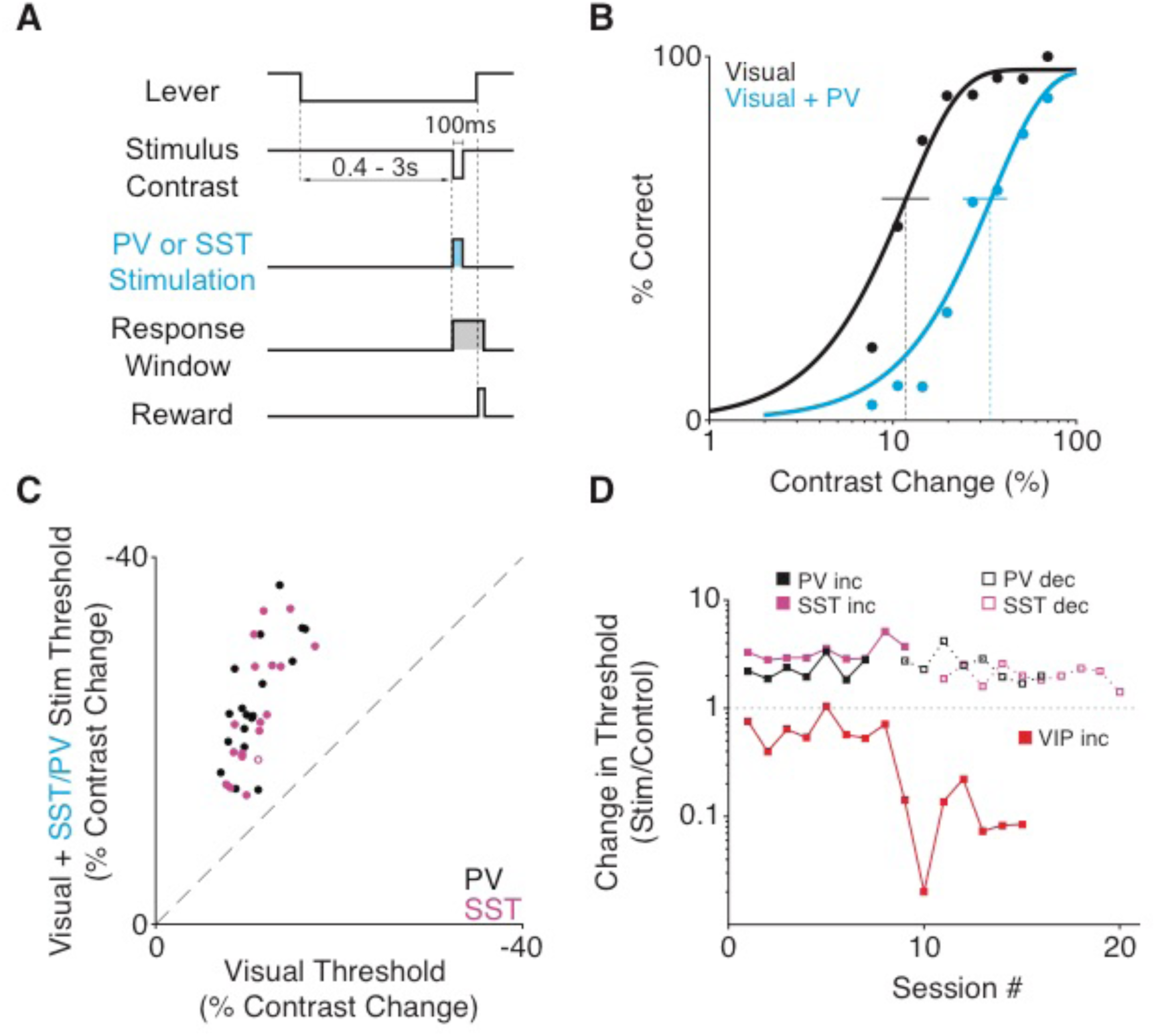
PV or SST stimulation impairs contrast decrement detection. (A) Trial schematic of the contrast decrement detection task. The visual stimulus contrast is fixed at 75% except during a 100 ms decrement. ChR2-expressing interneurons were illuminated with blue light for 100 ms concurrent with the contrast decrement on a randomly selected half of the trials. (B) Representative single session performance in the contrast decrement task for a PV mouse. Dots represent false-alarm corrected performance for trials with (blue) and without (black) activation of PV interneurons. Curves are best-fitting Weibull functions that were used to determine detection thresholds (dotted vertical lines) and 95% confidence intervals (solid horizontal lines). (C) Summary of PV and SST stimulation effects in the contrast decrement task. Circles depict the contrast decrement detection threshold from individual sessions with (y-axis) and without (x-axis) PV (black, 2 mice, 20 sessions) or SST (magenta, 2 mice, 18 sessions) stimulation. Filled circles represent sessions with a significant shift in threshold (37/38; bootstrapped). (D) Data from three representative mice showing that extended training does little to mitigate the impairment induced by PV (black) or SST (magenta) stimulation, while VIP (red) performance improves with training. Squares depict the ratio of thresholds from visual + optogenetic stimulus trials relative to visual only trials for each session. Values greater than one represent a perceptual impairment and values less than one represent performance improvement. Transition from solid to dashed lines in PV and SST mice depicts the switch from contrast increment to contrast decrement task.

In a follow up experiment, we tested whether the impairment in contrast decrement thresholds depended on the power used for optogenetic stimulation (n = 1 PV mouse). Our goal was to investigate whether increasing the intensity of inhibitory stimulation could produce a perceptible change in cortical output. We used different optogenetic powers (0.1 - 2.0 mW) in each session and measured the effect on performance (Figure S2). We found no evidence that different stimulation powers produced qualitatively different effects than those observed during the main experiments. The perceptual impairments scaled directly with the strength of inhibitory drive, falling almost linearly to zero as the optogenetic stimulation power was lowered (r^2^ = 0.67; Figure S2B).

These data suggest that animals cannot perceive the activation of PV or SST interneurons (or any change in activity they might cause in other neurons) and use that signal to guide a detection response. To determine if animals learned to detect interneuron activity, we examined optogenetically-induced changes in threshold across multiple sessions. Figure 3D plots threshold changes across training for representative mice. During first contrast increment (solid lines) and then contrast decrement tasks (dotted lines), stimulation of PV or SST neurons always impaired perception (threshold changes greater than one). This result was representative of all PV and SST mice (n=8). Across 125 total sessions (both increment and decrement), PV or SST stimulation never enhanced thresholds (threshold ratios less than one), indicating that stimulation never improved performance. The slope of a line fit to the example data from the PV mouse was not significantly different from zero (p = 0.96; F-test for linear regression). The slope of a line fit to the data from the SST mouse was significantly negative (p < 0.01; F-test for linear regression) indicating that threshold was elevated less in later sessions. However, only 1 of 5 PV mice and 1 of 3 SST mice showed a significant reduction in the threshold impairment. This suggests that any changes in the magnitude of impairment did not result from animals learning to detect PV or SST activation and instead could arise from behavioral strategies like discounting responses from affected neurons or reductions in perturbation strength via loss of ChR2 expressing neurons over time.

### Mice can learn to reliably report isolated VIP interneuron stimulation

In contrast to the largely stable impairments due to PV or SST activation, improvements in thresholds resulting from VIP stimulation grew over sessions (3D, red line). The change in the threshold ratio depended entirely on thresholds with VIP stimulation as performance on visual-only trials were similar early versus late in training (median = 6.5% versus 6.4; p = 0.66; Wilcoxon rank sum test). This indicates that mice can learn to use the activation of VIP neurons to guide their behavioral responses. Indeed, 2 of 3 VIP mice showed clear signs of learning, as performance on low contrasts (where the added optogenetic stimulus can make the largest contribution to performance) elevated across days (Figure S3). These data indicate that mice can learn to use VIP activity to guide their responses.

To examine this directly, we tested whether mice could be trained to respond to VIP neuron stimulation in the absence of natural visual stimulation. By progressively lowering the contrast of the visual stimulus, animals were conditioned to respond a 100-ms activation of VIP interneurons in isolation (Figure 4A). During each session, we varied the intensity of isolated VIP stimulation and measured the detection threshold for two mice (Figure 4B-C). Threshold estimates were relatively consistent across sessions (Figure 4D; mouse 1 mean threshold 0.09 mW, range 0.07–0.11 mW; mouse 2: mean threshold = 0.10 mW, Range 0.08–0.14 mW). Thus, the activity of VIP neurons produces changes in neuronal activity in sensory cortices that are perceptible and these changes can be used to guide behavioral responses.

**Figure 4.**
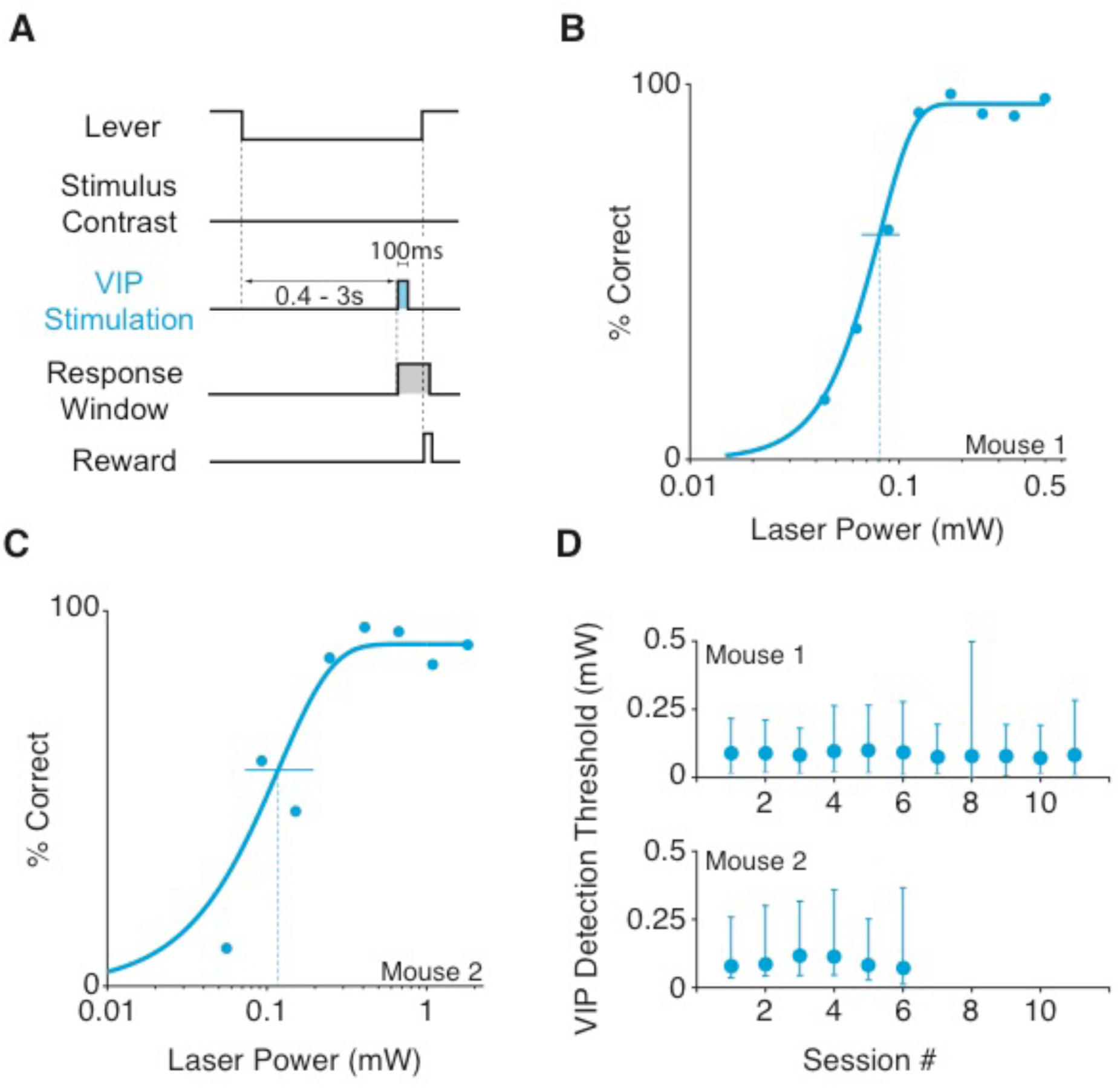
Mice can reliably report optogenetic stimulation of VIP neurons in the absence of a visual stimulus. (A) Trial schematic of the optogenetic stimulation detection task. The visual display contained a blank gray screen throughout the session. At a random time 400 to 3000 ms after trial onset a 100 ms square pulse of blue light was delivered to ChR2-expressing VIP neurons and the mouse was required to release the lever within the reaction time window to receive reward. (B) Representative psychometric performance of optogenetic stimulation detection in a VIP mouse. Dots represent false-alarm corrected performance as the intensity of VIP stimulation varied across trials. The curve is the best-fitting Weibull function, which was used to determine detection threshold (dotted vertical line) and 95% confidence interval (solid horizontal line). (C) Same as in B for a second VIP mouse. (D) Thresholds are stable across sessions. Top: VIP detection thresholds (95% CI) for 11 sessions from the mouse shown in B. Bottom: VIP detection thresholds (95% CI) for 6 sessions from the mouse shown in C.

## Discussion

We have shown that optogenetic activation of different interneuron classes produces distinct effects on visual contrast perception. Contrast increment detection was impaired by stimulation of either PV or SST interneurons, but improved by stimulation of VIP cells (Figure 2). Notably, activating either PV or SST neurons also impaired detection of contrast decrements, although enhanced suppression of pyramidal responses could have been expected to improve performance (Figure 3). Lastly, we found that mice could reliably detect optogenetic stimulation of VIP neurons in the absence of natural visual stimulation (Figure 4). We used transgenic mice to restrict expression of optogenetic actuators to genetically defined cell classes. However, even within a class, there exists substantial heterogeneity (Markram et al., 2004; Rudy et al., 2011). SST neurons exhibit layer dependent differences in activity (Muñoz et al., 2017) and diverse electrophysiological and morphological properties within a cortical layer (Maximiliano José et al., 2018). Similar differences have been reported for VIP interneurons (Prönneke et al., 2015). Thus, genetic labels may not capture the full diversity of interneuron effects on contrast perception. Nevertheless, our results provide new insights into how distinct sources of inhibition influence visual perception and highlight the types of neuronal signals that produce perceptible patterns of activity in downstream circuits.

### Comparisons with Previous Work

While multiple labs have investigated how PV, SST and VIP stimulation affects visual sensory representations (Adesnik et al., 2012; Atallah et al., 2012; Cottam et al., 2013; Fu et al., 2014; Lee et al., 2012; Pi et al., 2013; Wilson et al., 2012) relatively few studies have examined their role in perception, and with differing results.

In all prior work, PV neurons were stimulated before, during and after the onset of the target stimulus. However, dynamic inhibitory signals play a role in the perception of specific sensory events. We sought to study the effects of dynamic inhibitory responses that occur coincident with stimuli by restricting optogenetic perturbations to the brief stimulus epoch. Our data indicate that suppressing sensory responses concurrently with the onset of the visual stimulus can affect perception. Our behavioral results are supported by prior work examining the role of PV neurons in visual and somatosensory cortex (Glickfeld et al., 2013; Guo et al., 2014). These studies included neurophysiological recordings that showed reduced stimulus-evoked responses at the light intensities that affected behavior, suggesting the impairments in perceptual performance resulted from reduced spike output from sensory cortices.

A different study found that optogenetically enhancing PV activity in mouse V1 can improve orientation discrimination (Lee et al., 2012). The authors attributed this improvement to a sharpening of V1 tuning curves by PV stimulation. However, orientation discrimination thresholds were not measured. Instead the discriminability of specific, fixed orientation differences was assessed using a *d’* measure, which can improve significantly if perturbations cause the animal to withhold responses on a fraction of all trials. Earlier work from our lab showed that optogenetic stimulation of PV neurons impaired orientation discrimination thresholds (Glickfeld et al., 2013), suggesting that when mice are perceptually limited on orientation discrimination, PV-mediated effects in V1 do not improve orientation discrimination. This is supported by work in barrel cortex, where optogenetic activation of PV neurons impairs object location discrimination (Guo et al., 2014).

Another study found that optogenetically activating PV neurons in auditory cortex improved the ability of mice to detect a target stimulus presented on a background tone (Aizenberg et al., 2015). Interestingly, stimulus-evoked neuronal responses (relative to baseline firing rates) were enhanced by PV stimulation and this enhancement was correlated with the perceptual improvement across animals (Aizenberg et al., 2015). While potentiation of stimulus-evoked responses by enhanced inhibition is counterintuitive, simulated networks with high-gain excitatory units and strong stabilizing inhibitory feedback (Inhibition Stabilized Networks, or ISNs; Sadeh et al., 2017; Tsodyks et al., 1997) can exhibit this behavior following perturbations of the inhibitory population. Experimental evidence suggests that both visual and auditory cortices operate in an ISN-like regime (Kato et al., 2017; Ozeki et al., 2009). The findings of Aizenberg and colleagues might differ from our results because the more sustained photostimulation used in their study moved the cortical network to a different response regime. Future experiments could reconcile these differences.

Discrepancies such as these highlight challenges in using circuit perturbations to understand inhibitory neuron function. Strong optogenetic activation or suppression can produce new equilibrium states in cortex, not unlike those observed in ablation studies, that complicate interpretations of inhibitory neuron function (Sadeh et al., 2017). Furthermore, interdependent network interactions might mask the true nature of inhibition (Seybold et al., 2015). The choice of opsin also plays a critical role. Optogenetically exciting or inhibiting interneurons can produce asymmetrical effects on sensory responses (Phillips and Hasenstaub, 2016). Several opsins support optical inhibition (Wiegert et al., 2017) but can produce unintended effects on cellular activity or neurotransmitter release (Mahn et al., 2016; Raimondo et al., 2012), whereas different excitatory opsins produce a large range of photocurrents even with equivalent levels of expression (Klapoetke et al., 2014).

### Relating Changes in Neuronal Output to Behavior

A straightforward approach for detecting stimulus increments is to integrate stimulus-evoked activity to a decision criterion (Geisler, 2011). Indeed, we previously showed that detection of V1 activity was well described by linear integration of inputs over short intervals (Histed and Maunsell, 2014). Thus, the subtraction of spikes by PV or SST stimulation is a parsimonious explanation for impaired contrast increment detection. While there remains debate about whether PV or SST inhibition acts in a subtractive or divisive manner (Atallah et al., 2012; Lee et al., 2012; Seybold et al., 2015; Wilson et al., 2012), we found that the perceptual impairments induced by PV or SST stimulation were similar. This result is consistent with a recent data indicating that optogenetic stimulation of either PV or SST neurons reduces sensory responses in a predominantly linear fashion (Phillips and Hasenstaub, 2016).

It follows that detection of stimulus decrements could be accomplished by withholding responses until the spike rate falls below a decision criterion. If so, trials with PV or SST stimulation would produce a larger than normal decrement in neuronal activity and therefore enhance task performance. However, this notion is not supported by our data, as PV and SST stimulation impaired contrast decrement detection, even at high stimulation intensities (Figure S2). Our data raise the possibility that decrements in stimulus contrast are not decoded from decrements in V1 spiking. It has previously been suggested that retinal OFF pathways exists to convert stimulus decrements into an excitatory signal (Schiller, 1992). Opposed, rectified signalling channels can provide greater sensitivity and reduced latencies for spiking systems. It is possible that brain structures downstream of V1 decode stimulus increments and decrements based solely on the activity of neurons that increase their firing in response to those changes. This intriguing possibility might be addressed by detailed experiments examining correlations between population spiking patterns and particular perceptual outcomes.

VIP neurons primarily suppress the activity of SST neurons (Pfeffer et al., 2013), which amplify visual responses (Fu et al., 2014). Given that SST neurons primarily inhibit dendrites, activating VIP neurons could promote visual contrast detection by gating stimulus-related input onto the dendrites of V1 pyramidal neurons. However, we show that activity arising from VIP interneurons can be reliably perceived in the absence of visual stimuli. Thus, the performance improvement we observed could result from a distinct VIP-induced percept that the animals begin using to guide their responses and become more sensitive to with training. Alternatively, VIP-mediated improvements could be a combination of these processes. Given that optogenetic stimulation of V1 pyramidal neurons is perceptible (Histed and Maunsell, 2014) and VIP stimulation increases pyramidal cell spiking (Pi et al., 2013), VIP stimulation is likely to be detectable by virtue of activating V1 pyramidal neurons that project to other structures. Thus, even though VIP neurons comprise a small proportion of all cortical inhibitory neurons (Rudy et al., 2011), their influence on V1 spiking is sufficient to generate visual percepts.

## Methods

### Mouse Strains

All animal procedures were in compliance with the guidelines of the NIH and were approved by the Institutional Animal Care and Use Committee at The University of Chicago. Mouse lines were obtained from The Jackson Laboratory. Data come from parvalbumin-Cre mice (PV, 5 mice, 3 female; JAX stock #017320; Hippenmeyer et al., 2005), somatostatin-Cre mice (SST, 3 mice, 1 female; Jax stock #013044; Taniguchi et al., 2011), and vasointestinal peptide-Cre mice (VIP, 3 mice, 1 female, Jax stock #010908; Taniguchi et al., 2011) experimental animals were heterozygous for Cre recombinase in the cell type of interest (outbred by crossing homozygous Cre-expressing strains with wild type BALB/c mice, Jax stock #000651). Mice were singly housed on a reverse light/dark cycle with ad libitum access to food. Mice were water scheduled throughout experiments, except for periods around surgeries.

### Cranial window implant

Mice (3–5 months old) were implanted with a headpost and cranial window to give stable optical access for photostimulation during behavior (Glickfeld et al., 2013; Goldey et al., 2014; Histed and Maunsell, 2014). Animals were anesthetized with ketamine (40 mg/kg, i.p.), xylazine (2 mg/kg, i.p.) and isoflurane (1.2– 2% in 100% O_2_). Using aseptic technique, a headpost was secured using C&B Metabond (Parkell) and a 3 mm craniotomy was made over the left cerebral hemisphere (3 mm lateral and 0.5 mm anterior to lambda) to implant a glass window (0.8 mm thickness; Tower Optical).

### Intrinsic autofluorescence imaging

We located V1 by measuring changes in the intrinsic autofluorescence signal using epifluorescence imaging (Andermann et al., 2011). Autofluorescence produced by blue excitation (470 ± 40 nm, Chroma) was collected using a green long-pass filter (500 nm cutoff) and a 1.0x air objective (Zeiss; StereoDiscovery V8 microscope; ∼0.11 NA). Fluorescence was captured with a CCD camera (AxioCam MRm, Zeiss; 460×344 pixels; 4×3 mm field of view) For retinotopic mapping; we presented full contrast sinusoidal drifting gratings (10° SD Gabor patch; 30°/s; 0.1 cycles/deg) for 10 s followed by 6 s of mean luminance. The response to the visual stimulus was computed as the fractional change in fluorescence during the first 8 s of the stimulus presentation compared to the average of the last 4 s of the preceding blank.

### Viral injections and ChR2 stimulation

ChR2 injections were targeted to a monocular region of V1 based on each animal’s retinotopic map (+25° in azimuth; between −15° to +15° in elevation). Before virus injection, mice were anesthetized (isoflurane, 1–1.5%), and the glass window was removed using aseptic technique. We used a volume injection system (World Precision Instruments) to inject 200 nl of AAV9-Flex-ChR2-tdTomato (∼10^11^ viral particles; Penn Vector Core) 300 µm below the pial surface. The virus was injected at a rate of 40 nl/min through a glass capillary attached to a 10 μL syringe (Hamilton). Following the injection, a new cranial window was sealed in place. Several weeks after injection, we localized the area of ChR2 expression using tdTomato fluorescence, and attached an optical fiber (400 µm diameter; 0.48 nA; Doric Lenses) within 500 µm of the cranial window (∼1.3 mm above the cortex). We delivered light though the fiber from a 455 nm LED (ThorLabs) and calibrated the total power at the entrance to the cannula. Optogenetic stimulation began no earlier than 6 weeks after injection. We prevented optogenetic stimuli from cueing the animal to respond by wrapping the fiber implant in blackout fabric (Thor Labs) that secured to the headpost using a custom mount.

### Behavioral tasks

Animals were water scheduled and trained to respond to changes in a visual display using a lever (Histed et al., 2012). Animals were first trained to respond to the appearance of a Gabor stimulus on uniform background with the same average luminance. The Gabor stimulus (SD 6.75°, 0.1 cycles/deg, odd-symmetric) appeared for 100 ms and its contrast varied randomly from trial to trial across a range that spanned behavioral threshold. Stimuli for each animal were positioned at a location that corresponded to the V1 representation expressing ChR2 (25° azimuth, −15° to +15°). Mice initiated trials by depressing a lever. After a random delay (400-3000 ms), the Gabor appeared for 100 ms, and the mouse had to release the lever within a brief response period beginning 100 ms after stimulus onset (max: 900 ms). Early releases and misses resulted in a brief timeout before the start of the next trial. Behavioral control and data collection and analysis were done using custom software based on MWorks (http://mworks-project.org), Matlab (MathWorks) and Python.

Optogenetic stimulation did not begin until animals worked reliably for hundreds of trials each day and contrast detection thresholds were stable across days. This typically required ∼2 months of training. During optogenetic experiments, we activated ChR2 expressing inhibitory neurons on a randomly selected half of the trials by delivering blue light through the optical fiber positioned over V1. Opsin illumination was always concurrent with the visual stimulus and at a fixed power (adjusted initially to cause an approximately two-fold change in detection threshold).

Following data collection using the contrast increment task, a subset of PV (2 mice; 1 female) and SST (2 mice, both male) were trained (without optogenetic stimulation) to perform a contrast decrement detection task. The timing and response parameters remained the same, but a 75% contrast Gabor was on the display except during a randomly timed 100 ms interval, when it decremented to one of a set of pre-assigned contrasts. The mouse had to release the lever within the response window to receive a reward. With most of the mice, performance stabilized in the contrast decrement task within ∼7 sessions, and we began optogenetic stimulation of the same patch of ChR2 expressing interneurons that were stimulated during increment sessions. As with the contrast increment task, optogenetic stimulation occurred on a randomly selected half of trials at a fixed power.

Some VIP mice were to report changes in neuronal activity induced by ChR2 in the absence of a visual stimulus. In these experiments, the video display was set to a mid-level gray through each session. Optogenetic stimulation was presented for 100 ms at a randomly selected time during each trial, and at a power that was randomly selected from a pre-assigned set of values.

### Histology

Mice were perfused with 10% pH-neutral buffered formalin (Millipore Sigma Inc.), after which the brain was removed and submerged in fixative for 24 hr. The brain was subsequently rinsed with PBS, placed in a 30% sucrose PBS solution until it sank. Brains were sectioned at 40 µm on a freezing microtome, mounted and cover slipped. tdTomato expression was visualized with 561 nm excitation using a Caliber I.D. RS-G4 Confocal Microscope with a 10x objective (Olympus; 0.3 NA).

### Data analysis

Sessions in which the false alarm rate was greater than 50% or the mouse missed more than 50% of trials were excluded from analysis. Detection thresholds were determined using trials in which the subject either responded correctly (hit) or failed to respond (miss, Histed et al., 2012). We corrected for false alarms by finding the rate of false alarms in each 100 ms bin during trials. We subtracted a fraction of correct trials from each contrast level based on the time of responses and the false alarm rate function. This correction was typically small (median hits removed 6.9%; range, 2.3–13.0% for 168 sessions from 11 mice). Corrected performance data were then fit with Weibull cumulative distribution functions using non-linear least squares and variance weighting of each mean. The two psychometric functions (with and without ChR2 stimulation) were fit using four parameters: individual thresholds, a common lapse rate, and a common slope. A small fraction of sessions (24/168) were better fit by the addition of a second slope parameter specific to ChR2 stimulation trials (F-Test for number of parameters). Sessions fit with 5 parameters were distributed across genotypes (PV: 10.4%; SST: 14.7%; VIP: 16.2% of total sessions) and produced similar effects of stimulation on psychometric detection thresholds (median change in threshold (Visual + Optogenetic Threshold/ Visual Threshold) with 4 Parameters = 1.74; median threshold change with 5 parameters = 1.56; p = 0.50; Wilcoxon rank-sum test). Threshold confidence intervals were estimated using a bootstrap (1000 repetitions, p < 0.05, one-tailed). In the case of ChR2 detection in the absence of visual stimuli (VIP-Cre mice), we fit a three parameter Weibull function (lapse rate, slope, threshold). False alarm rates in optogenetic detection experiments were comparable to experiments with both visual and optogenetic stimuli (percentage of trials removed; median, 6.8%; range, 5.8–12.8%; *n* = 17 sessions from 2 mice.

## Supplemental Figures and Legends

**Figure S1.**
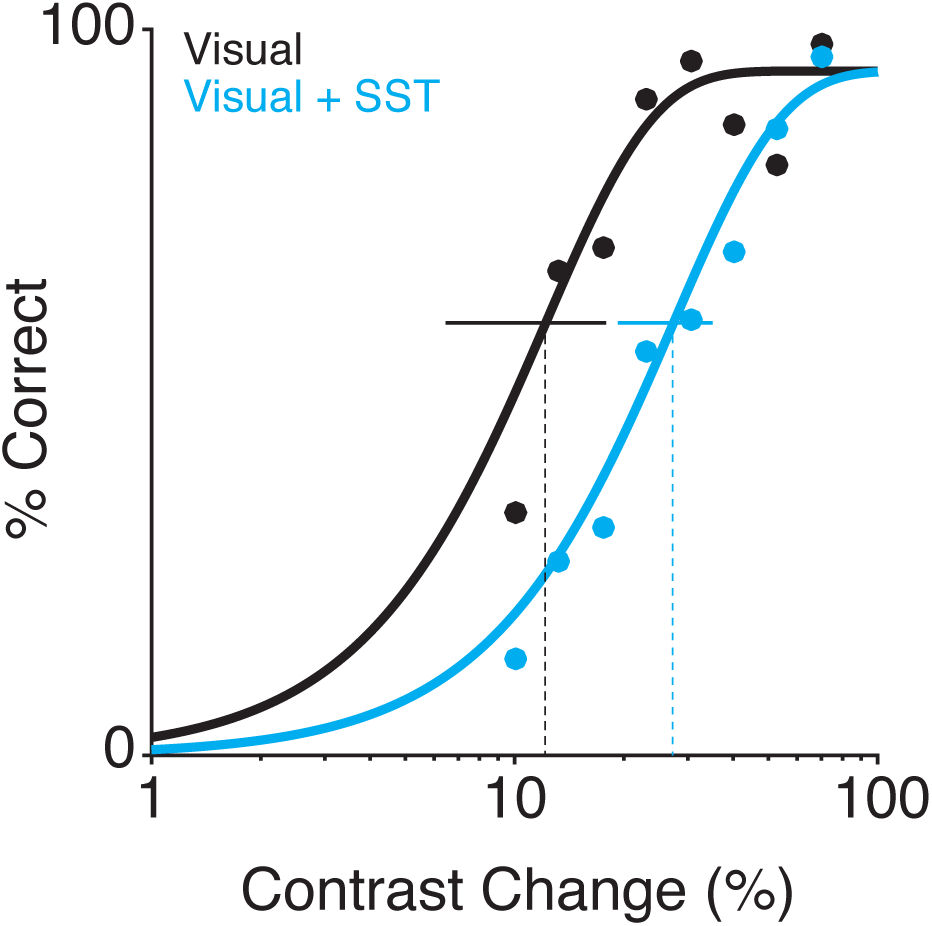
Activation of SST neurons impairs performance in the contrast decrement detection task. Representative behavioral performance from a single contrast decrement session in an SST mouse. Filled dots represent false-alarm corrected performance (% correct) for trials with (blue) and without (black) activation of SST interneurons as the magnitude of contrast decrement is varied. Data are fit with a Weibull function to determine detection thresholds (dotted vertical lines) and 95% confidence intervals (solid horizontal lines).

**Figure S2.**
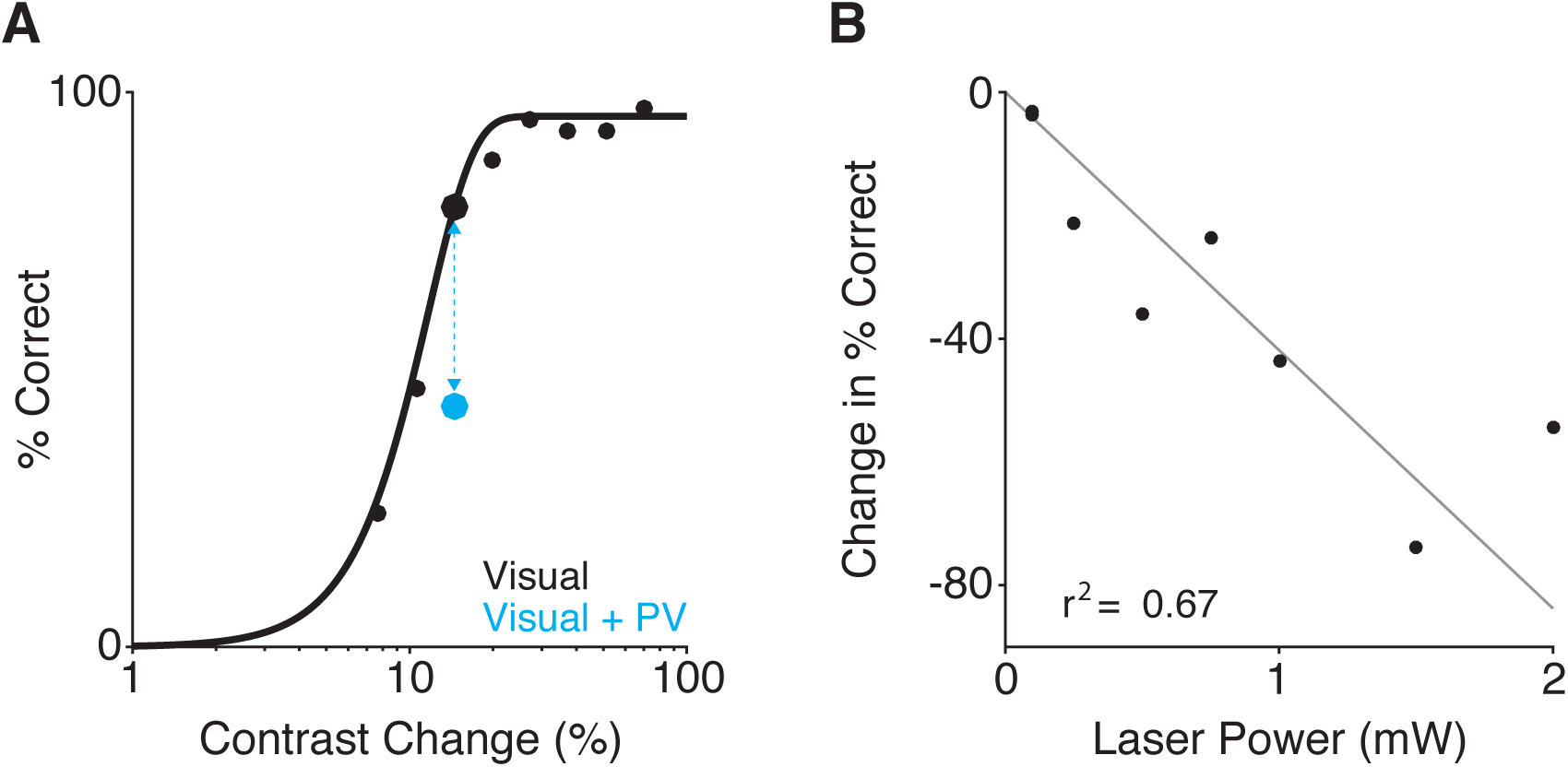
Magnitude of perceptual impairment induced by PV stimulation scales with stimulation intensity. (A) Representative behavioral performance from a single contrast decrement session where one visual contrast decrement was randomly paired with stimulation of PV interneurons. Optogenetic power of 0.5 mW was the stimulation intensity used in this example session (and also the power used for this mouse throughout the main experiment). Filled dots represent performance for trials with (blue) and without (black) activation of PV interneurons as the magnitude of contrast decrement is varied. Solid line depicts performance on visual only trials fit with a Weibull function. Blue dashed line connecting dots shows magnitude of perceptual impairment on the selected contrast decrement with (blue) without (black) optogenetic activation of PV neurons. (B) The perceptual impairment induced by PV activation scales with optogenetic stimulation intensity. Individual points depict performance impairment from individual sessions (note: there are two sessions at 0.1 mW). Gray line is a maximum likelihood linear regression anchored at the origin (r^2^ = 0.67).

**Figure S3.**
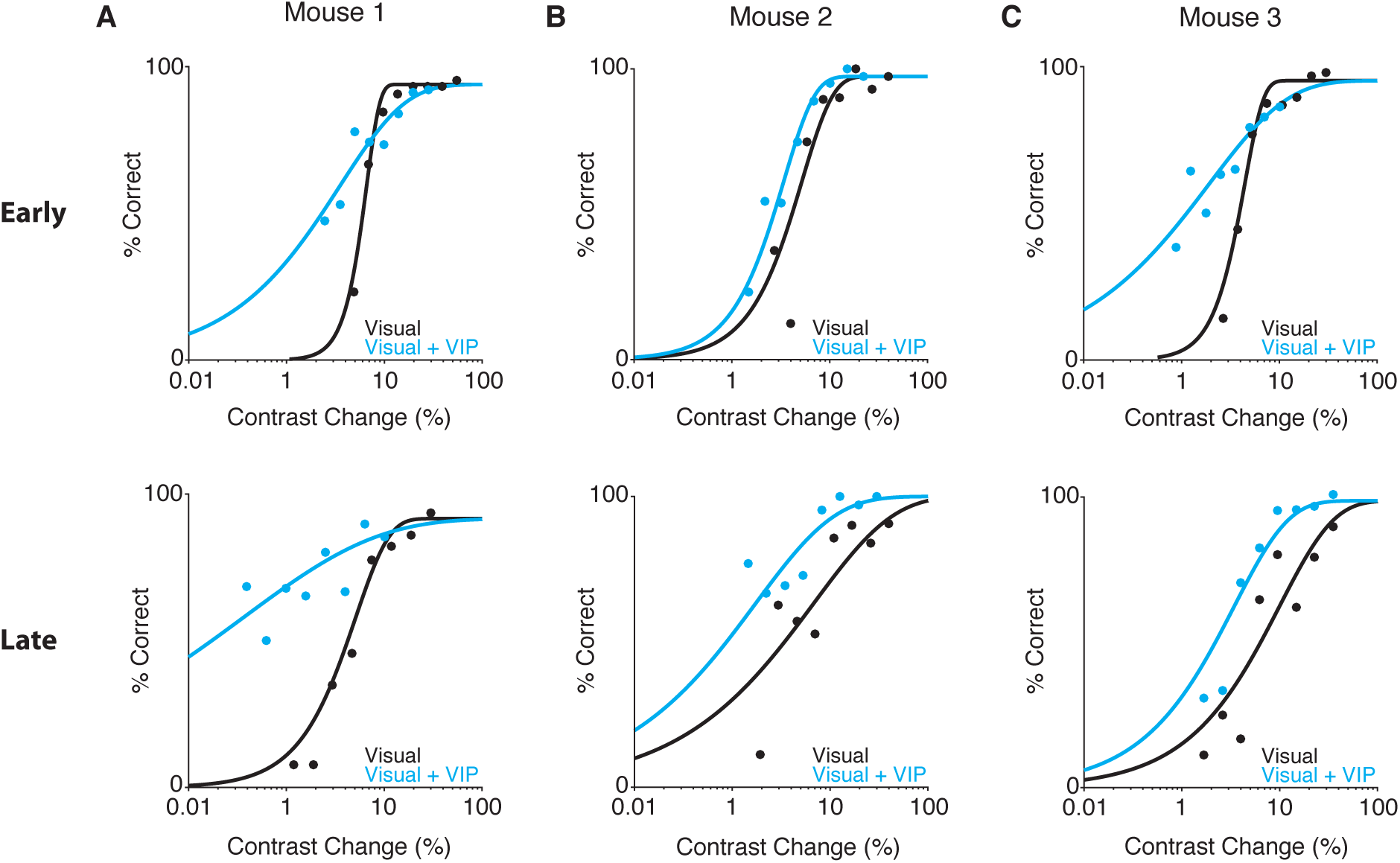
VIP mice improve on responses to VIP stimulation across sessions. Representative psychometric functions measured early (top row) versus late (bottom row) in training for the 3 VIP mice (columns) used in this study. All sessions are separated by at least 5 days of training during which the stimulation intensity was held constant. Filled dots represent performance for trials with (blue) without (black) activation of VIP interneurons as the contrast increment is varied. (A) Example sessions from mouse 1. With training, performance at visual contrast steps that were too small to be reliably detected improved noticeably. (B) Similar results were obtained in mouse 2, as performance improved over time for low contrast trials in the presence of VIP stimulation. (C). Data from mouse 3, whose performance did not improve after extended training.

**Table S1.** Summary of performance data from sessions with concurrent visual and optogenetic input. Inc. = contrast increment sessions. Dec = contrast decrement sessions.

|  | **PV+ Inc.** | **SST+ Inc.** | **VIP+ Inc.** | **PV+ Dec.** | **SST+ Dec.** |
| --- | --- | --- | --- | --- | --- |
| Mice | 5 (3F, 2M) | 3 (1F, 2M) | 3 (1F, 2M) | 2 (1F, 1M) | 2 (2M) |
| Sessions | 47 | 40 | 43 | 20 | 18 |
| Sessions w.<br>Significant<br>Threshold Shift | 44 (93.6%) | 40 (85%) | 43 (93%) | 20 (100%) | 17 (94.4%) |
| Change<br>in Threshold<br>Range | 1.0 - 4.3 | 1.1 - 5.1 | 0.02 - 1.0 | 1.3 - 4.2 | 1.3 - 3.0 |
| Sessions Fit<br>With 5<br>Parameters | 5 (10.6%) | 7 (17.5%) | 7 (16.2%) | 2 (10%) | 3 (16.7%) |
| False Alarm<br>Rate | 6.3% | 5.9% | 7.8% | 6.8% | 8.3% |
| False Alarm<br>Rate Range | 2.3% - 11% | 3.0% - 12% | 5.9% - 11% | 5.4% - 8.8% | 4.5% - 13% |

**Table S2.**
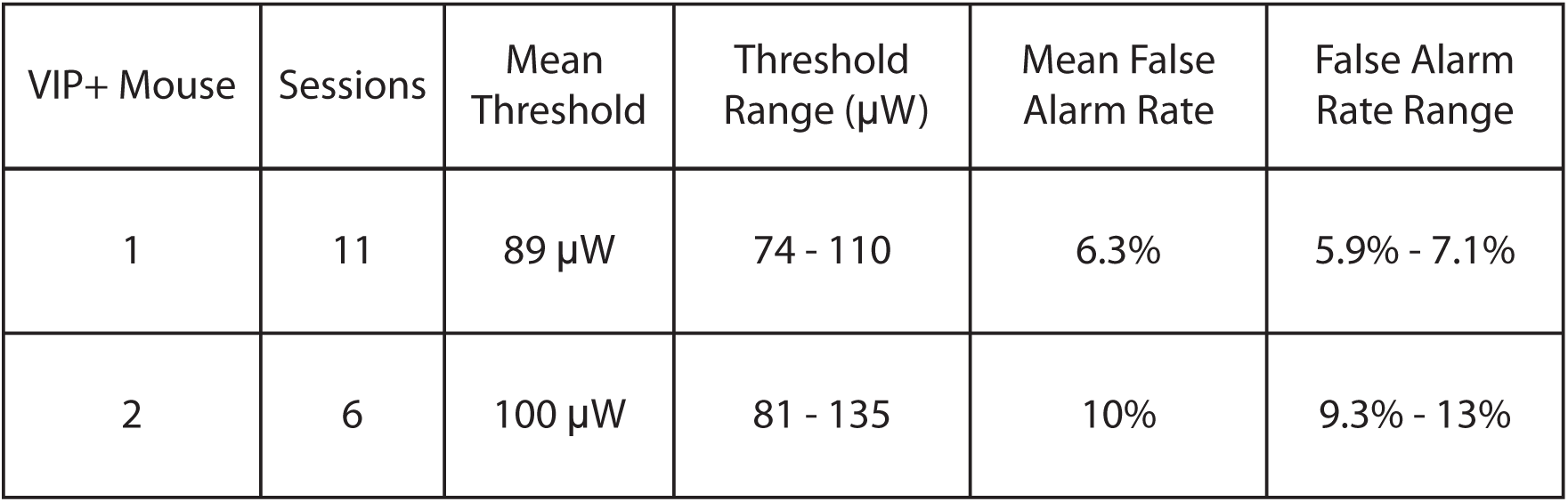
Summary of performance data from VIP optogenetic detection sessions

**Author Contributions**
JJC, MHH, and JHRM designed research. JJC and MDS conducted research. JJC analyzed data. JJC and JHRM wrote the manuscript with input from MHH and MDS.

## Acknowledgements

The authors thank Dr. Vytas Bindokas and the University of Chicago Integrated Light Microscopy Core Facility for assistance with confocal imaging and Elizabeth Page for assistance with animal training. This work was supported by an Albert O. Beckman Postdoctoral Fellowship (JJC) and NIH Grant U01-NS090576.

## References

Adesnik, H., Bruns, W., Taniguchi, H., Huang, Z.J., and Scanziani, M. (2012). A neural circuit for spatial summation in visual cortex. Nature 490, 226–230.

Aizenberg, M., Mwilambwe-Tshilobo, L., Briguglio, J.J., Natan, R.G., and Geffen, M.N. (2015). Bidirectional Regulation of Innate and Learned Behaviors That Rely on Frequency Discrimination by Cortical Inhibitory Neurons. PLoS Biol. 13, 1–32.

Andermann, M.L., Kerlin, A.M., Roumis, D.K., Glickfeld, L.L., and Reid, R.C. (2011). Functional specialization of mouse higher visual cortical areas. Neuron 72, 1025–1039.

Atallah, B. V., Bruns, W., Carandini, M., and Scanziani, M. (2012). Parvalbumin-Expressing Interneurons Linearly Transform Cortical Responses to Visual Stimuli. Neuron 73, 159–170.

Carandini, M., and Heeger, D.J. (2012). Normalization as a canonical neural computation. Nat. Rev. Neurosci. 13, 51–62.

Cottam, J.C.H., Smith, S.L., and Hausser, M. (2013). Target-Specific Effects of Somatostatin-Expressing Interneurons on Neocortical Visual Processing. J. Neurosci. 33, 19567–19578.

Ferster, D. (1988). Spatially opponent excitation and inhibition in simple cells of the cat visual cortex. J. Neurosci. 8, 1172–1180.

Fu, Y., Tucciarone, J.M., Espinosa, J.S., Sheng, N., Darcy, D.P., Nicoll, R.A., Huang, Z.J., and Stryker, M.P. (2014). A cortical circuit for gain control by behavioral state. Cell 156, 1139–1152.

Geisler, W.S. (2011). Contributions of ideal observer theory to vision research. Vision Res. 51, 771–781.

Glickfeld, L.L., Histed, M.H., and Maunsell, J.H.R. (2013). Mouse primary visual cortex is used to detect both orientation and contrast changes. J. Neurosci. 33, 19416–19422.

Goldey, G.J., Roumis, D.K., Glickfeld, L.L., Kerlin, A.M., Reid, R.C., Bonin, V., and Andermann, M.L. (2014). Versatile cranial window strategies for long-term two-photon imaging in awake mice. Nat. Protoc. 9, 2515–2538.

Guo, Z.V., Li, N., Huber, D., Ophir, E., Gutnisky, D., Ting, J.T., Feng, G., and Svoboda, K. (2014). Flow of Cortical Activity Underlying a Tactile Decision in Mice. Neuron 81, 179–194.

Hippenmeyer, S., Vrieseling, E., Sigrist, M., Portmann, T., Laengle, C., Ladle, D.R., and Arber, S. (2005). A developmental switch in the response of DRG neurons to ETS transcription factor signaling. PLoS Biol. 3, 0878–0890.

Histed, M.H., and Maunsell, J.H.R. (2014). Cortical neural populations can guide behavior by integrating inputs linearly, independent of synchrony. Proc. Natl. Acad. Sci. U. S. A. 111, E178–87.

Histed, M.H., Carvalho, L.A., and Maunsell, J.H.R. (2012). Psychophysical measurement of contrast sensitivity in the behaving mouse. J. Neurophysiol. 107, 758–765.

Kato, H.K., Asinof, S.K., and Isaacson, J.S. (2017). Network-Level Control of Frequency Tuning in Auditory Cortex. Neuron 95, 412–423.e4.

Klapoetke, N.C., Murata, Y., Kim, S.S., Pulver, S.R., Birdsey-Benson, A., Cho, Y.K., Morimoto, T.K., Chuong, A.S., Carpenter, E.J., Tian, Z., et al. (2014). Independent optical excitation of distinct neural populations. Nat. Methods 11, 338–346.

Kubota, Y., Karube, F., Nomura, M., and Kawaguchi, Y. (2016). The Diversity of Cortical Inhibitory Synapses. Front. Neural Circuits 10, 1–15.

Lee, S.H., Kwan, A.C., Zhang, S., Phoumthipphavong, V., Flannery, J.G., Masmanidis, S.C., Taniguchi, H., Huang, Z.J., Zhang, F., Boyden, E.S., et al. (2012). Activation of specific interneurons improves V1 feature selectivity and visual perception. Nature 488, 379–383.

B. Liu, hua, Y. Li, tang, W. Ma, pei, C. Pan, jie, Zhang, L.I., and Tao, H.W. (2011). Broad inhibition sharpens orientation selectivity by expanding input dynamic range in mouse simple cells. Neuron 71, 542–554.

Mahn, M., Prigge, M., Ron, S., Levy, R., and Yizhar, O. (2016). Biophysical constraints of optogenetic inhibition at presynaptic terminals. Nat. Neurosci. 19, 554–556.

Markram, H., Toledo-Rodriguez, M., Wang, Y., Gupta, A., Silberberg, G., and Wu, C. (2004). Interneurons of the neocortical inhibitory system. Nat. Rev. Neurosci. 5, 793–807.

Maximiliano José, N., Hashikawa, Y., and Rudy, B. (2018). Diversity and connectivity of layer 5 somatostatin-expressing interneurons in the mouse barrel cortex. J. Neurosci. 2415–2417.

Muñoz, W., Tremblay, R., Levenstein, D., and Rudy, B. (2017). Layer-specific modulation of neocortical dendritic inhibition during active wakefulness. Science (80-.). 355, 954 LP–959.

Nagel, G., Szellas, T., Huhn, W., Kateriya, S., Adeishvili, N., Berthold, P., Ollig, D., Hegemann, P., and Bamberg, E. (2003). Channelrhodopsin-2, a directly light-gated cation-selective membrane channel. Proc. Natl. Acad. Sci. 100, 13940–13945.

Ozeki, H., Finn, I.M., Schaffer, E.S., Miller, K.D., and Ferster, D. (2009). Inhibitory Stabilization of the Cortical Network Underlies Visual Surround Suppression. Neuron 62, 578–592.

Pfeffer, C.K., Xue, M., He, M., Huang, Z.J., and Scanziani, M. (2013). Inhibition of inhibition in visual cortex: The logic of connections between molecularly distinct interneurons. Nat. Neurosci. 16, 1068–1076.

Phillips, E.A.K., and Hasenstaub, A.R. (2016). Asymmetric effects of activating and inactivating cortical interneurons. Elife 5, 1–22.

Pi, H.J., Hangya, B., Kvitsiani, D., Sanders, J.I., Huang, Z.J., and Kepecs, A. (2013). Cortical interneurons that specialize in disinhibitory control. Nature 503, 521–524.

Priebe, N.J., and Ferster, D. (2008). Inhibition, Spike Threshold, and Stimulus Selectivity in Primary Visual Cortex. Neuron 57, 482–497.

Prönneke, A., Scheuer, B., Wagener, R.J., Möck, M., Witte, M., and Staiger, J.F. (2015). Characterizing VIP neurons in the barrel cortex of VIPcre/tdTomato mice reveals layer-specific differences. Cereb. Cortex 25, 4854–4868.

Raimondo, J. V., Kay, L., Ellender, T.J., and Akerman, C.J. (2012). Optogenetic silencing strategies differ in their effects on inhibitory synaptic transmission. Nat. Neurosci. 15, 1102–1104.

Rudy, B., Fishell, G., Lee, S.H., and Hjerling-Leffler, J. (2011). Three groups of interneurons account for nearly 100% of neocortical GABAergic neurons. Dev. Neurobiol. 71, 45–61.

Sadeh, S., Silver, R.A., Mrsic-Flogel, T.D., and Muir, D.R. (2017). Assessing the Role of Inhibition in Stabilizing Neocortical Networks Requires Large-Scale Perturbation of the Inhibitory Population. J. Neurosci. 37, 12050–12067.

Schiller, P.H. (1992). The ON and OFF channels of the visual system. Trends Neurosci. 15, 86–92.

Seybold, B.A., Phillips, E.A.K., Schreiner, C.E., and Hasenstaub, A.R. (2015). Inhibitory Actions Unified by Network Integration. Neuron 87, 1181–1192.

Taniguchi, H., He, M., Wu, P., Kim, S., Paik, R., Sugino, K., Kvitsani, D., Fu, Y., Lu, J., Lin, Y., et al. (2011). A Resource of Cre Driver Lines for Genetic Targeting of GABAergic Neurons in Cerebral Cortex. Neuron 71, 995–1013.

Tsodyks, M., Skaggs, W.E., Sejnowski, T.J., and Mcnaughton, B.L. (1997). Paradoxical effects of inhibitory interneurons. J. Neurosci. 17, 4382–4388.

Wiegert, J.S., Mahn, M., Prigge, M., Printz, Y., and Yizhar, O. (2017). Silencing Neurons: Tools, Applications, and Experimental Constraints. Neuron 95, 504–529.

Wilson, N.R., Runyan, C.A., Wang, F.L., and Sur, M. (2012). Division and subtraction by distinct cortical inhibitory networks in vivo. Nature 488, 343–348.

